# Gestational exposure to bisphenol S induces microvesicular steatosis by promoting lipogenesis and inflammation in male rat offspring

**DOI:** 10.1101/2023.06.01.543354

**Authors:** Archana Molangiri, Saikanth Varma, Kota Sri Naga Hridayanka, Myadara Srinivas, Suryam Reddy Kona, Ahamed Ibrahim, Asim K Duttaroy, Sanjay Basak

## Abstract

Fetal exposure to endocrine-disrupting bisphenol A (BPA) showed a long-lasting programming effect on organ development and predisposed to the metabolic risk of adult diseases. However, limited data on developmental exposure to BPA-substitute bisphenol S (BPS) in predisposing liver metabolic disease is available. Here, the effects of BPS exposure were assessed on hepatic metabolism by examining adiposity and inflammation in the adipose and liver of the 90-day male offspring. Pregnant Wistar rats were exposed to BPA and BPS (0.0, 0.4, 4.0 µg/kg bw) via gavage from gestational day 4 to 21. Prenatal BPS-exposed offspring exhibited a higher obesogenic effect than BPA, including changes in body weight, body fat, feed efficiency, and leptin signalling. The fasting blood glucose did not change, but BPS exposure elevated plasma corticosterone levels and adipocyte hypertrophy of the visceral adipose tissue (VAT) to a greater extent than BPA. Adipocyte hypertrophy was augmented by modulated expression of lipid uptake (PPARγ, FABP4), glucocorticoid (HSD11β1), inflammation (IL6, IL1β, CRP, COX2), oxidative stress (CHOP) and apoptotic (Caspase 3) mediators. Liver histology showed numerous lipid droplets, and hepatocyte ballooning, associated with upregulated expression of cholesterol, lipid biogenesis and glucocorticoid activators, indicating microvesicular steatosis in the prenatally BPS-exposed adult offspring. The upregulated PPARα, ADRP, and FGF21 expression and increased lipid peroxidation in the offspring’s liver suggest metaflammation due to fetal exposure to BPS. Fetal BPS exposure demonstrated a more significant disruption in metabolism involving adiposity, liver fat, inflammation in excess, and predisposition to hepatic steatosis in the male offspring.

**Highlights:**

- Fetal BPS exposure exhibited enlarged and inflamed adipocytes more than BPA
- Prenatal BPS exposure induced excess lipid droplets & hepatocyte ballooning in liver
- In utero exposure to BPS induces microvesicular steatosis in adult rats

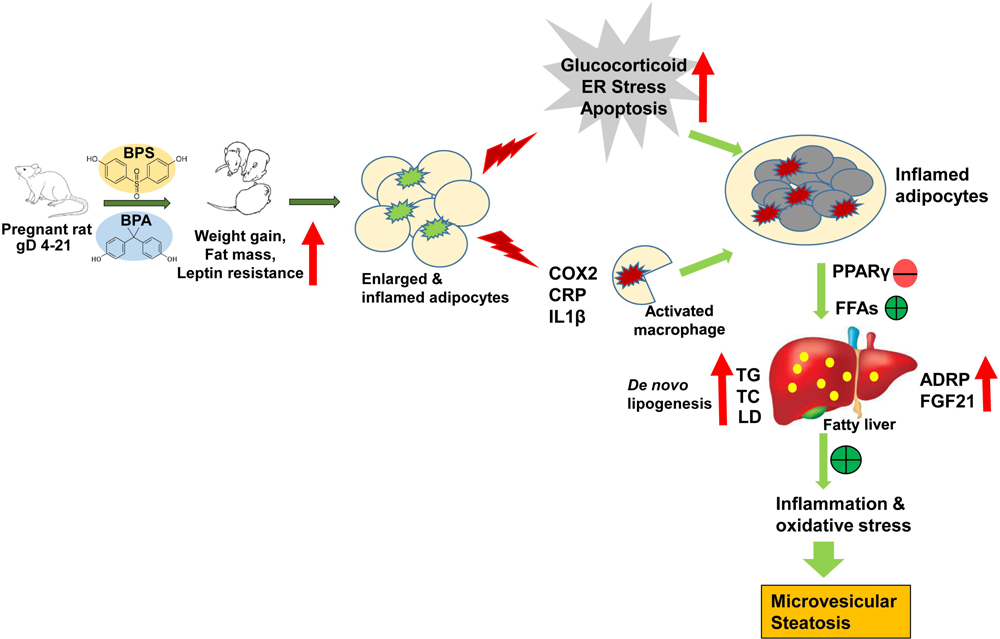

## 1. Introduction

The environmental endocrine disrupting chemical (EDC) such as bisphenol A (BPA) is ubiquitously exposed to all (Vandenberg et al., 2010), from dermal to oral, as well as by atmospheric route (Kassotis et al., 2015). Due to the potential health risk of exposure to BPA [4,4-(propane-2,2-diyl) diphenol], the industry replaced it by substituting the central carbon atom with two methyl and carbonyl groups [4,4′-dihydroxydiphenyl sulphone], as bisphenol S (BPS). The usage of “BPA-free” products mainly contain BPS (Moon, 2019), which is documented by human exposure to this contaminant (Jacobson et al., 2019), with much higher systemic retention than BPA (Khmiri et al., 2020), thus poses new biological risks. The exposure window is critical, particularly in early life during setting up organ function that could determine disease risk later (Basak et al., 2020). Prenatal exposure to EDC mixtures, including phenolic substances, is associated with liver injury risk and susceptibility to developing liver metabolic disease later (Midya et al., 2022) and contributes to developing hepatic steatosis (Treviño and Katz, 2017). Urinary BPA is linked with liver abnormalities (Lang et al., 2008), and urinary BPS is associated with increased obesity and abdominal obesity (Jacobson et al., 2019). Higher exposure to BPS and BPA increases the incidence of simple hepatic steatosis due to elevated serum cholesterol (An et al., 2021; Peng et al., 2023). Long-term BPS exposure triggered the development of hepatic steatosis by increased triacylglycerol (TAG) accumulation (Wang et al., 2019) and modulating lipid metabolism, endoplasmic reticulum (ER) stress (Qin et al., 2020) in male zebrafish. Numerous pathways reveal that adipose-liver crosstalk contributes to the progression and development of fatty liver disease (Cimini et al., 2017). Inflammation in adipose tissue (AT) secretes free fatty acids (FFAs) into the systemic circulation and production of pro-inflammatory cytokines leading to secondary inflammation in the liver, indicating close interplay among adipose tissue and liver (Yang et al., 1997).

Several *in vivo* studies suggest that BPA exposure affects an individual’s metabolic profile by altering the lipid profile, and lipolytic gene expression leads to obesity (Naomi et al., 2022). In contrast, excessive glucocorticoid exposure may lead to central adiposity and dyslipidemia (Wake and Walker, 2004). The enzyme HSD11β1 (11β-hydroxysteroid dehydrogenase type 1) rejuvenates active corticosterone in rodents by activating and multiplying the glucocorticoid receptor, which promotes preadipocyte differentiation to adipose hypertrophy. Prenatal BPA exposure led to the escalation of adipocyte hypertrophy, over-expressing of peroxisome proliferator-activated receptor gamma (PPARγ) and increased lipogenesis in the liver (Somm et al., 2009), accelerated the development of hyperlipidemia and obesity by altering serum leptin (Miyawaki et al., 2007). Promoting hepatic steatosis by modifying *de novo* fatty acid synthesis and altering lipogenic genes are suggestive outcomes of BPA-contaminated foods (Marmugi et al., 2012). Exposure to BPA leads to hepatotoxicity involving edema, hepatocellular decay, and necrosis (Khan et al., 2016). Gestational BPA exposure dysregulated adipogenesis & lipid homeostasis (Fang et al., 2022), and induced the development of fatty liver (Long et al., 2021) in the offspring. Moreover, BPA exposure from pre-gestation through lactation in combination with a high-fat diet (HFD) produced hepatic steatosis in dams (Marchlewicz et al., 2022).

BPA analogues have an equal impact on the obesity (Andújar et al., 2019). Concerns arise since BPA-alternative-BPS exhibits perpetual effects upon its exposure (Chen et al., 2016). The estimated plasma clearance of BPS in humans is two times lower than BPA, indicating a more significant exposure risk of replacing BPA with BPS (Gayrard et al., 2020). BPS exhibited almost identical or greater metabolic effects in several in vivo studies (Oliviero et al., 2022). For example, BPS exposure builds up obesity by interfering with glucose metabolism and lipid accumulation in the liver, triggering inflammations (Meng et al., 2019). Prenatal BPS exposure promotes inflammation in the offspring’s reproductive tissue by upregulating IL6 and apoptosis (Molangiri et al., 2022). Fetal BPS exposure results in overweight male mice by inducing excess lipid storage in the fat tissue (Ivry Del Moral et al., 2016). Prenatally BPS-exposed adult male mice exhibited adipose hypertrophy, expansion in adipocyte size, and PPARγ activation and were predisposed to HFD-induced adipogenesis (Ahn et al., 2020). Gestational BPS exposure affects terminal differentiation gene expression of preadipocytes (Pu et al., 2017) and modulates pre-adipocyte differentiation and metabolic signalling by binding to the PPARγ ligand (Schaffert et al., 2021).

The participation of several mediators, including PPARs, fatty acid-binding protein 4 (FABP4), and adipose differentiation-related protein (ADRP) in adipose-liver crosstalk, contributes to the progression of fatty liver disease. BPA exposure induced PPARγ activation by modulating the lipid metabolic gene expression and dysregulated metabolism (Gao et al., 2020). PPARγ and PPARα are predominantly expressed in various tissues and modulates adipogenesis and liver metabolism. FABP4, a lipid trafficking chaperone protein, is the primary target for PPARγ corresponding to obesity-associated anomalies, postulated as a causal factor in fatty liver progression (Moreno-Vedia et al., 2022). FABP4 predominantly resides in the adipocytes, whose expression led to the upregulation of lipogenesis via PPARγ modulation due to BPA exposure (Gao et al., 2020). FABP4 enhances adipogenesis by downregulating PPARγ, working vice versa (Garin-Shkolnik et al., 2014), altering metabolic events leading to inflammation. BPA exposure promoted lipid droplet formation, ROS accumulation in the liver (Song et al., 2019), and upregulated lipid droplet protein (ADRP) expression in the male gonads (Varma et al., 2023).

The present study investigates the endocrine effects of prenatal BPA and BPS exposure on adipose-liver histology, interaction with liver-specific PPARα and its regulation on lipid droplet protein, ADRP, and differentiation growth factor, FGF21, in the offspring liver.

## 2. Material and methods

### 2.1 Animal experiment

In brief, 3-months female Wistar rats (n=40) and body weight-matched male rats (n=20) were selected. Rats were housed in two per cage and kept on a 12 h light/dark cycle at 22°C ± 2°C, with a relative humidity of 45-55%. They were given a standard chow diet for 7 days and water *ad libitum*. The estrus phase confirmed the fertile window. Mating was initiated (Females: males = 2:1), and conception was confirmed by the vaginal plug discharge and considered a gestational day (gD 0). Bisphenols (BPA and BPS) were gavaged from gD 4 until gD 21, as described before (Molangiri et al., 2022). Exposure was avoided during the initial 4 days to prevent implantation failure. The control (olive oil) did not receive bisphenols. The other pregnant rats received bisphenols of 0.4, 4.0 μg/kg body weight (bw) per day, considering the tolerable daily intake of BPA exposure by the European Food Safety Authority. The offspring continued with a standard chow diet. The data were collected from male rats from 30 to 90 days of the offspring’s age. The study design and procedures (#ICMR-NIN/IAEC/02/008/2019) were approved by the Institutional Animal Ethical Committee of ICMR-National Institute of Nutrition, which conformed to the guidelines of the Care and Use of Laboratory Animals.

### 2.2 Assortment of blood and tissues

Fasting blood was collected by puncturing the retro-orbital plexus using an antiseptic capillary tube. Plasma and serum were removed and stored at −80°C for subsequent analysis. Carbon dioxide asphyxiation was employed to euthanize 90 days rats to collect liver and adipose. The organ and tissue weight were recorded and stored separately in 4% PBS buffered formalin for histology, RNA later, and snap-frozen in liquid nitrogen for mRNA and protein expression respectively.

### 2.3 Food intake, body weight, and dual-energy X-ray absorptiometry

Body weights of the rat offspring were determined fortnightly, starting from their age of 45 days. The rats were gently restrained, and their body weights were documented on an electronic weighing balance. Daily food intake was measured for two consecutive weeks from 45 days onward. The rats were fed with ∼ 30 g of chow pellet diet per day and estimated the food intake by considering the leftover. For the body fat composition, the rats were anaesthetized with a combination of ketamine & xylazine (80mg/kg and 10mg/kg bw), delivered intraperitoneally. A complete body scan was performed on DEXA (Discovery, Hologic, Bedford, MA, USA), which reported body mass, lean body mass, fat mass, and fat percentage.

### 2.4 Biochemical assays

The fasting glucose, triacylglycerol, and cholesterol (total) were measured in the plasma (Biosystems, Barcelona, Spain). Hepatic triacylglycerol and cholesterol were also measured in liver tissue. Tissue homogenate was prepared in 5% aqueous Nonidet P40 (NP40).

### 2.5 Plasma cortisol by immunoassay

The plasma cortisol was measured by ELISA (Fine test, Cat # ER1651). Briefly, 50 µl of plasma, standards, and biotin-labelled antibodies were loaded onto the pre-coated plate, followed by incubation at 37°C for 45 min, washed, and HRP-streptavidin conjugate was added. Color was developed by TMB substrate and measured at 450 nm in a microplate reader (Bio-Tek, Powerwave XS) after the addition of a stop reagent. Concentrations are expressed as ng/ml.

### 2.6 Tissue histology by hematoxylin and eosin (H&E) staining

The tissues (visceral adipose and liver) were stored in 10% formalin and embedded in the paraffin blocks. The tissue sections (∼ 4μm) were prepared using a microtome. The sections were deparaffinized, rehydrated, and stained in hematoxylin solution for 20-40 min. As the stain developed, the slides were immersed in 70% ethanol containing 1% HCl (acid ethanol), which led to the removal of excess dye. The slides were subsequently stained with Eosin for 10 min and washed in 100% ethanol for 5 min. Finally, the slides were rinsed in xylene, mounted with a cover slip to avoid any air bubbles trapped inside, and dried overnight.

### 2.7 Immunofluorescence

The liver tissues were embedded in paraffin blocks, and microsections (4 µm) were prepared on a glass slide. Deparaffinization, antigen retrieval, and non-specific blocking of slides were performed as described (Varma et al., 2023). The primary antibodies, such as Anti-ADRP, and anti-FGF21, were added to the slides and incubated overnight at 4°C. Goat anti-rabbit (Alexa fluorTM 488 #A-11034) was added against the primary antibody and mounted with DAPI medium (#F6057, Sigma, St. Louis, MI, USA). Fluorescence was captured in an inverted microscope (Nikon Eclipse TE2000-U).

### 2.8 Immunoblotting

The immunoblotting was performed and analyzed to measure the expression of proteins in tissues (visceral adipose and liver) as described previously (Molangiri et al., 2022), except that membrane (blot) were probed with different sets of primary antibodies (**Supplementary Table 1**).

### 2.9 Quantitative real-time-PCR (qRT-PCR)

The expression of various genes was measured by quantifying mRNA fold levels as described previously (Varma et al., 2023), except for a different set of SYBR green primers (Sigma Merck, **Supplementary Table 2**). The obtained Ct values were analyzed using the ddCt method for relative quantification of mRNA expression derived from three independent experiments, using the expression of β-actin mRNA as the endogenous control.

### 2.10 TBARS assay

Lipid peroxidation was used to evaluate oxidative stress in liver tissue. The assay was performed as described before ((Molangiri et al., 2022).

### 2.11 Statistical analysis

Multiple group comparison analysis was performed using One-way ANOVA in GraphPad Prism v.8. platform. Every experiment was performed independently and repeated multiple times, as mentioned in the text or figure legends. Data were considered statistically significant when the p-value fell less than 0.05. The data were represented as mean ± standard error of the mean (SEM).

## 3. Results

### 3.1 Exposure to BPS during pregnancy increased the offspring’s body weight, similar to BPA, as evidenced by the longitudinal assessment

Compared to the control, the body weight of the offspring was increased significantly (p<0.05) with different concentrations of BPA and BPS exposed (0.4 and 4.0 μg/kg) when measured at 45d, 59d, and 73d of their ages (**Fig.1 A-C**). No significant difference was observed in the body weight gain between BPA and BPS-exposed rats. Multiple comparisons showed a significant time-depended increase in the body weight of the rat offspring over a while (45d to 73d).

**Fig.1.**
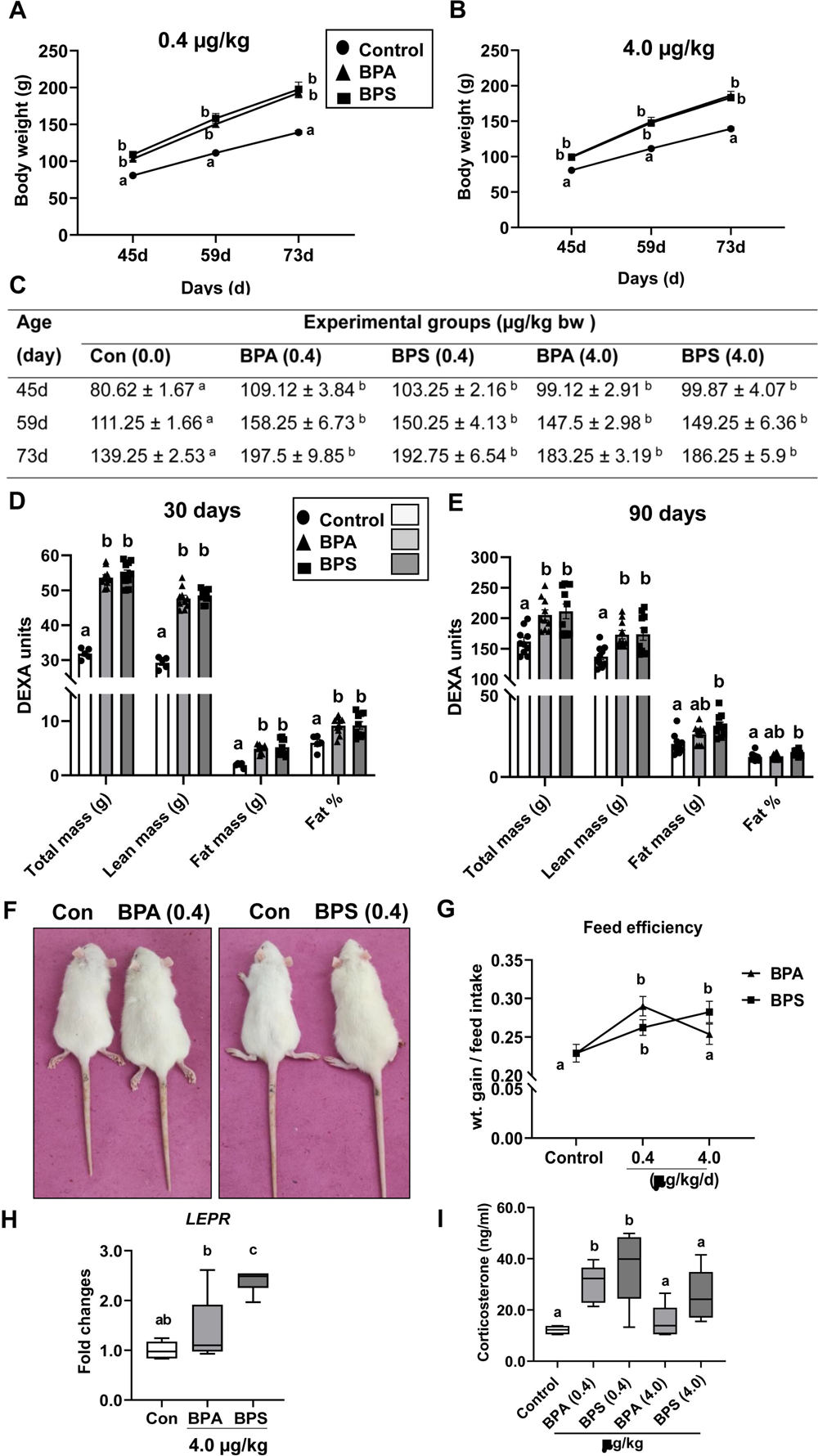
Body weight, body composition, feed efficiency, and leptin signalling mediator in the offspring following *in utero* bisphenols (BPA and BPS) exposure. (A-C) Body weight of the male offspring at 45, 59, and 73 days of age due to bisphenol exposure (A) 0.4µg/kg bw (B) 4.0 µg/kg. (C) Repeated measurement was performed by two-way ANOVA and expressed as mean ± SEM, n=8/group. (D-E) Dual-energy X-ray absorptiometry (DEXA) measured the body composition of the rat offspring at (D) 30 days (E) 90 days of their age, where dams were exposed to bisphenols (0.4µg/kg bw, n=10/group). (F) Representative images of 60 d offspring rats prenatally exposed to BPA and BPS (0.4µg/kg bw). (G) Feed efficiency was calculated based on weight gain (g) over feed intake (g) from 45-60 days (n=10/group). (H) Expression of leptin receptor (LEPR) mRNA in the offspring liver employing RT-qPCR after normalizing with endogenous control β-actin (n=6/group). (I) Plasma corticosterone (ng/ml) of 90 d male offspring rats exposed to bisphenols (n=6-8/group). Data were analyzed by one-way ANOVA with Tukey’s multiple comparison tests and expressed as mean ± SEM. Values with unlike superscript letters were significantly different p<0.05 vs control.

### 3.2 Fetal BPS exposure changed the body composition consistently as the offspring grew

Since the body weights were altered longitudinally, the body composition was measured at 30 and 90 d of age at the lowest BPA and BPS (0.4 μg/kg) exposure (**Fig.1 D-E**). Offspring (30d) showed a significant increase (p<0.05) in body mass, lean mass, fat mass, and fat percentage compared to the control. The increased body fat (fat mass and fat percentage) remained significant (p<0.05) only for BPS-exposed offspring in their adult (90d) age (**Fig.1E**). However, all bisphenol-exposed offspring (60d) acquired obese phenotype (**Fig.1F**). Their feed efficiency was considerably increased in a concentration-dependent manner in BPS-exposed offspring (**Fig.1G**). The LEPR expression was profoundly upregulated in the offspring liver, which was significantly (p<0.05) higher in BPS (**Fig.1H**) compared to BPA and control. The expression of other signalling mediators such as leptin (LEP), adiponectin (ADIPOQ), neuropeptide Y (NPY), and proopiomelanocortin (POMC), those control food intake, did not change among these groups (data not shown).

### 3.3 BPS-exposed offspring released higher corticosterone than BPA

Gestational bisphenols (BPA and BPS) exposure did not alter plasma glucose, total triacylglycerol (TAG), and total cholesterol (TC) in 90d offspring (**Supplementary Table 3**). In contrast, plasma corticosterone level was significantly increased (p<0.05) with 0.4µg/kg bisphenols (BPA and BPS) exposed adult offspring, and the trend (p>0.05) was continued with higher exposure (4.0 µg/kg). The BPS-exposed rats showed higher plasma corticosterone than BPA at comparable concentrations (**Fig1I**).

### 3.4 Prenatal BPS exposure induced adipocyte hypertrophy and inflammation in the offspring to a greater extent than BPA

Adipose histology exhibited irregularly shaped, disorganized adipocytes due to bisphenols exposure, possibly inflamed and enriched with hypertrophic adipocytes **(Fig.2A i-v)**. Prenatal BPS exposure significantly increased the mean adipocyte diameter to a greater extent than BPA in the dose-dependent manner [control vs. BPA and BPS (0.4μg/kg): 42.46±1.66 vs. 60.28±3.29 and 76.72±3.57; vs. BPA and BPS (4.0 μg/kg): 74.34±3.92 and 90.03±6.37 (µm), p<0.05, **Fig.2B**]. The expression of FABP4 was upregulated (p<0.05) in BPS-exposed offspring’s adipose tissue (**Fig.2C**), while PPARγ (**Fig.2D**) expression was (p<0.05) downregulated in these tissues. The expression of HSD11β mRNA, which converts inactive cortisone to the active form, was significantly increased in BPS (p<0.05, **Fig.2E**), while HSD11β2 expression was unaffected (data not presented). The HSD11β1 protein expression was (p<0.05) higher in bisphenol exposed (4.0 μg/kg) adipose tissue than in the control (**Fig.2F**).The expression of pro-inflammatory cytokines and ER stress markers was increased in the offspring’s adipose tissue. The expression of IL1β, CRP, and COX2 was significantly increased to a different degree (∼2 to 30 fold, **Fig.3 A-C**) and significantly increased CHOP (C/EBP homologous protein) and caspase 3 expression **(Fig.3 D-E)** indicating the presence of a large number of stressed apoptotic adipocytes in BPS-exposed adipose tissue.

**Fig.2.**
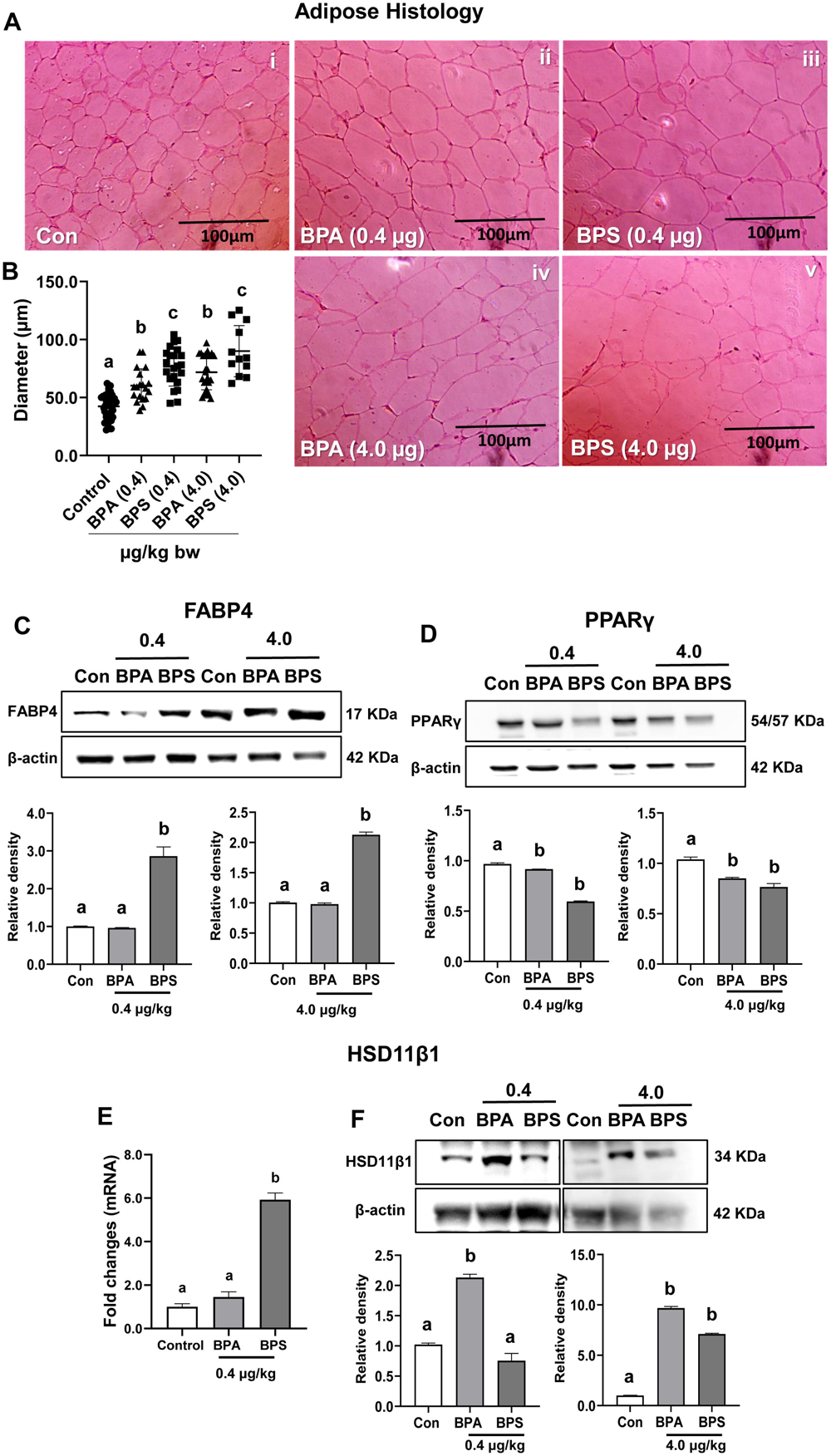
Adipose tissue (visceral) histology and expression of metabolic modulators of 90d offspring exposed to bisphenols *in utero*. (A) Representative sections show visceral adipocytes stained with haematoxylin and eosin (20X). (B) Adipocyte diameter is expressed in μm (n=15-20/group). Immunoblots and relative expression of (C) FABP4, and (D) PPARγ show the normalized protein expressions (n=3 rats/group). The normalized mRNA expression of (E) HSD11β1 employing RT-qPCR (n=6/group) and (F) protein expression of HSD11β1. Values are expressed as mean ± SEM, and the implication of unlike letters is considered statistically significant at p < 0.05 vs. control.

**Fig.3.**
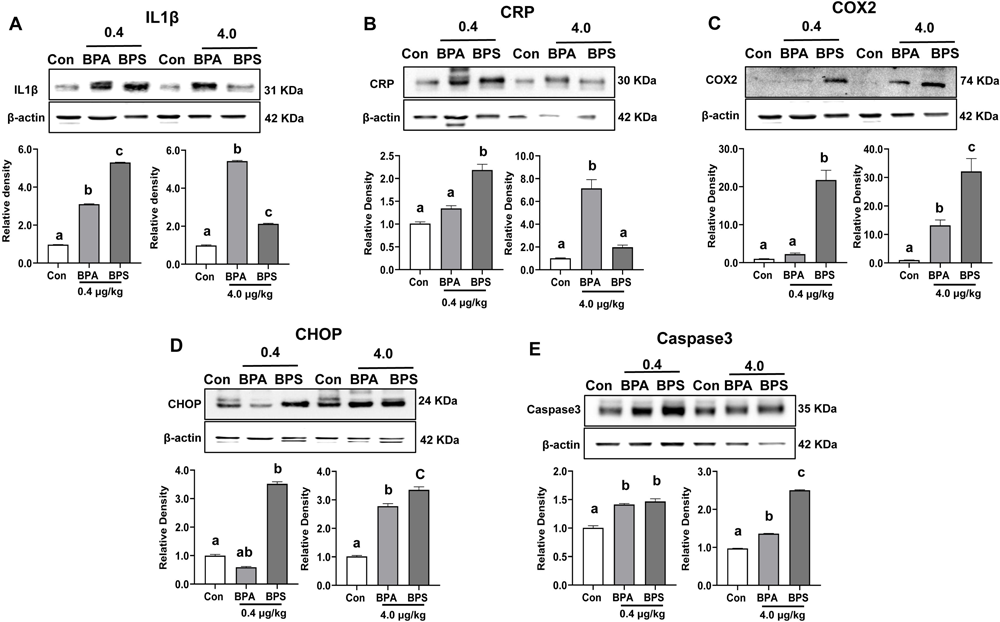
Expression of pro-inflammatory cytokines and ER stress-induced apoptosis mediators in the offspring’s adipose tissue (visceral) following prenatal bisphenol exposure. Immunoblots and the relative expression of (A) IL1β, (B) CRP, (C) COX2, (D) CHOP, and (E) Caspase 3 show the normalized protein expression over β-actins. Respective bars indicate the relative band intensity of normalized expressed proteins. Data represent mean ± SEM (n=3 rats/group) using one-way ANOVA with Tukey’s multiple comparison test. Values with unlike letters indicate significantly different at p<0.05 vs. control.

### 3.5 Prenatal BPS exposure induced microvesicular steatosis in the offspring

Liver phenotypes were altered in bisphenols-exposed offspring, including their colour and surface. Control livers appeared dark with a smooth texture. In contrast, BPS-exposed livers exhibited greater discolouration with a greasy surface and more changes in the liver lobes than in control, and BPA **(Fig.4A).** Liver weights were significantly increased due to BPS exposure [control vs BPA and BPS (0.4μg/kg): 6.58±0.24 vs. 10.84±0.44 and 11.22±0.51; vs BPA and BPS (4.0 μg/kg): 8.21±0.42 and 10.02±0.53 (g), p<0.05, **Fig.4B**]. Liver histology displayed a regular arrangement of hepatocytes in control rats (**Fig.4C i)**. Excess inter-hepatocyte fats, infiltrated neutrophils, and hepatocyte ballooning were noted in the bisphenol-exposed liver **(Fig.4C ii-v).** The clusters of azonal foamy hepatocytes as microvesicular patches were accompanied by medium-sized lipid droplets **(> 50%)** in BPS-exposed liver **(Fig.4C iii &v).** The distended or inflated hepatocytes, in which the nucleus is centrally located, indicated the predominant presence of microvesicular steatosis in the BPS-exposed offspring liver **(Fig.4C iii & v).** Although BPA-exposed liver also displayed lipid droplets **(<20%)** and microvesicular patches **(Fig.4C ii & iv)**, their intensity was less in numbers in comparison to BPS. The elevated levels of total cholesterol and triacylglycerol content in the liver homogenates **(Fig.4 D-E)** and fat droplets with ballooning hepatocytes in the BPS-exposed liver are the marked features of microvesicular steatosis in adult rats.

**Fig.4.**
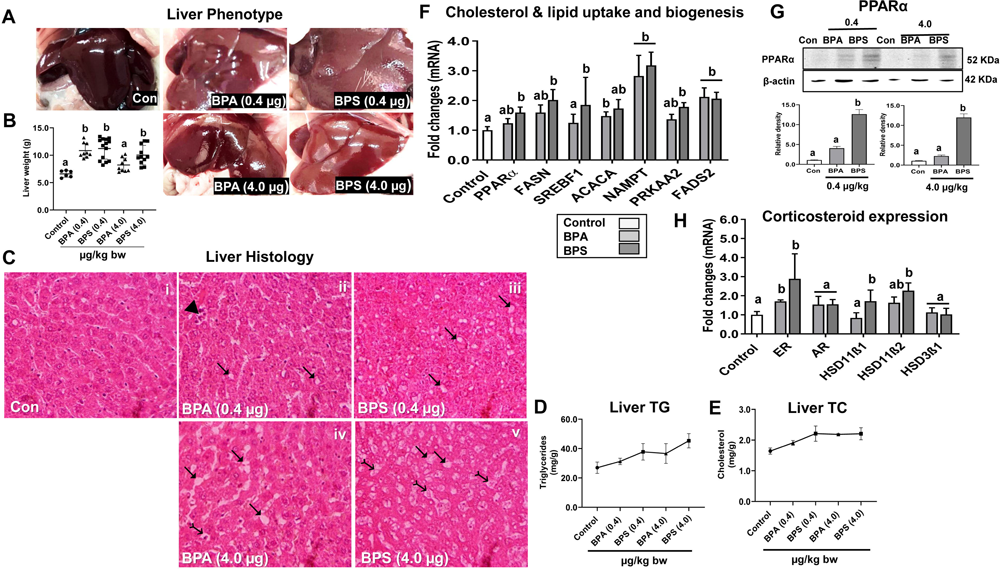
Macroscopic, microscopic, biochemical and molecular investigation of the 90d offspring liver exposed to bisphenols *in utero*. (A) Gross images displayed the offspring’s liver phenotypes. (B) The liver weight of the rat offspring (n=7-12/group). (C) Representative photomicrographs (40X) of liver histology, stained with H&E. Marked symbols are used on the images to define distinct aspects of the changes in liver histology. IZ pointing triangle represents small necrotic regions with focal aggregations of inflammatory cells. IZ triangle headed arrows indicate the presence of lipid droplets. ↣ Inward arrows pinpoint hepatocytes ballooning along with the centrally located nucleus, indicative of microvesicular steatosis corresponding to fatty liver/liver injury. (D) Total triglyceride (TG) and (E) total cholesterol (TC) levels in the offspring (90d) liver tissue (n=6/group). Expression of mRNAs associated with (F) cholesterol and lipid biogenesis and (H) steroidogenesis and glucocorticoid mediators. Expression levels were measured using RT-qPCR after normalising the expression of each mRNA with the expression of endogenous control β-actin. Fold changes of normalized mRNA expression of bisphenol groups are compared over the normalized expression of the control group (n=4-6/group). (G) Immunoblots of PPARα and quantitative bars indicating their relative expression in the liver (n=3 rats/group). Data were analysed with one-way ANOVA with Tukey’s multiple comparison test. Values are represented as means ± SEM. ^a, b^ values with unlike letters were significantly different vs. control at p<0.05.

### 3.6 Hepatic steatosis was augmented by upregulated expression of lipid biogenesis modulators and glucocorticoid mediators in offspring liver

Prenatal BPS exposure promoted *de novo* lipogenesis by significant (p<0.05) upregulation in the expression of PPARα, FASN, SREBF1, ACACA, NAMPT, PRKAA2 and FADS2 mRNAs in the liver comparable or more significant than BPA (**Fig.4F**), while mRNA expression of PPARγ, CEBPα, FABP4, IGF1, GLUT4, SCD1, FADS1, ELOVL2, ELVOL5 did not change (p>0.05) (data not presented). The PPARα expression was upregulated by more than ∼12 fold (p<0.05) only in BPS-exposed (0.4 and 4.0 μg/kg) liver (**Fig.4G**), while PPARγ expression was unaltered (data not presented). The expression of ER mRNA was significantly upregulated (p<0.05) in response to BPA and BPS exposure, with a higher magnitude in the BPS-exposed liver **(Fig.4H)**. The expression of HSD11β1 and HSD11β2 mRNAs were significantly upregulated exclusively in BPS-exposed liver **(Fig.4H)**.

### 3.7 In utero, bisphenol exposure increased ADRP & FGF21 protein accumulation and their expression in the offspring’s liver

A relatively higher ADRP expression (>5 fold) across the concentrations of BPS-exposed offspring indicated a potential increase in the lipid load of the BPS-exposed liver than BPA (**Fig.5A**). The CTCF of ADRP was significantly and dose-dependently increased in the BPS-exposed liver [control vs. 0.4 and 4.0 BPS (μg/kg): 0.17±0.03 vs. 1.56±0.35 and 2.34±0.19 (au), p<0.05, **Fig.5B-C**]. The increased FGF21 **(Fig.6A)** expression indicated an altered liver metabolism, possibly due to an imbalance between lipid oxidation and lipogenesis in the bisphenols-exposed offspring. The localized FGF21 expression was increased with bisphenol exposure [control vs. BPA and BPS (0.4μg/kg): 0.48±0.06 vs. 1.44±0.23 and 2.42±0.16; vs BPA and BPS (4.0 μg/kg): 4.82±0.39 and 3.61±0.39 (au), p<0.05, **Fig.6 B-C**].

**Fig.5.**
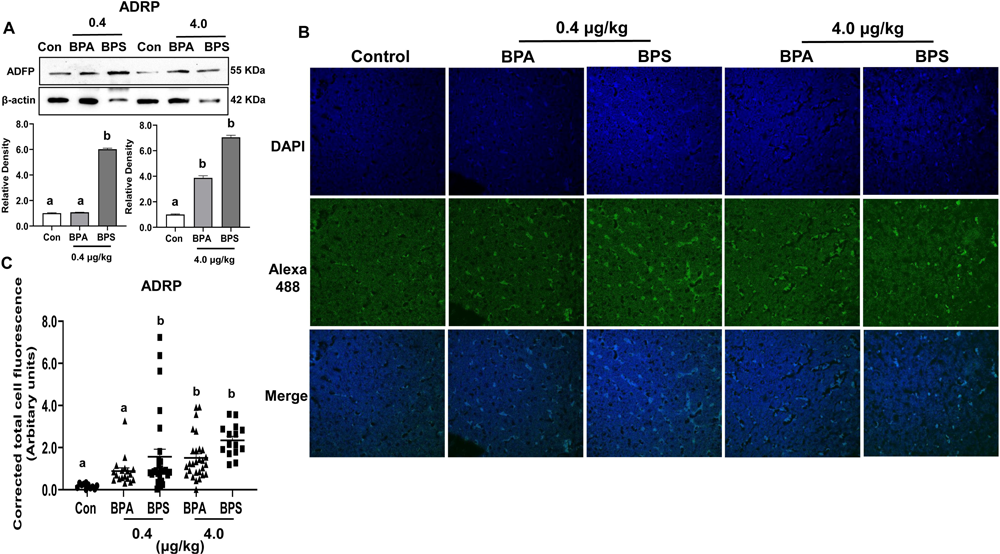
The expression and localization of lipid droplet protein, ADRP by immunofluorescence in the 90d offspring liver exposed to bisphenols prenatally. (A) Immunoblots of ADRP and its relative expression in the liver (n=3 rats/group). (B) Localized expression of ADRP by immunofluorescence is depicted in the representative images. Liver tissue was fixed in Bouin’s solution, dehydrated, and paraffinized. The staining of tissue sections was performed with Alexa Fluor 488 tagged-ADRP antibody (green) and nuclei with DAPI (blue). Images were captured with a confocal laser scanning microscope (Leica Microsystems, Germany) at 63X magnification using an oil immersion objective. (C) The total fluorescence was quantified using ImageJ software (NIH, USA). Corrected total cell fluorescence (CTCF) of ADRP expression was calculated after measuring the fluorescence [CTCF= integrated density – (area of selected cell x mean fluorescence of background readings)] and expressed in arbitrary units. Data were analyzed with one-way ANOVA with Tukey’s multiple comparison tests and expressed as means ± SEM (n=3). Values with unlike letters were significantly different vs control at p<0.05.

**Fig.6.**
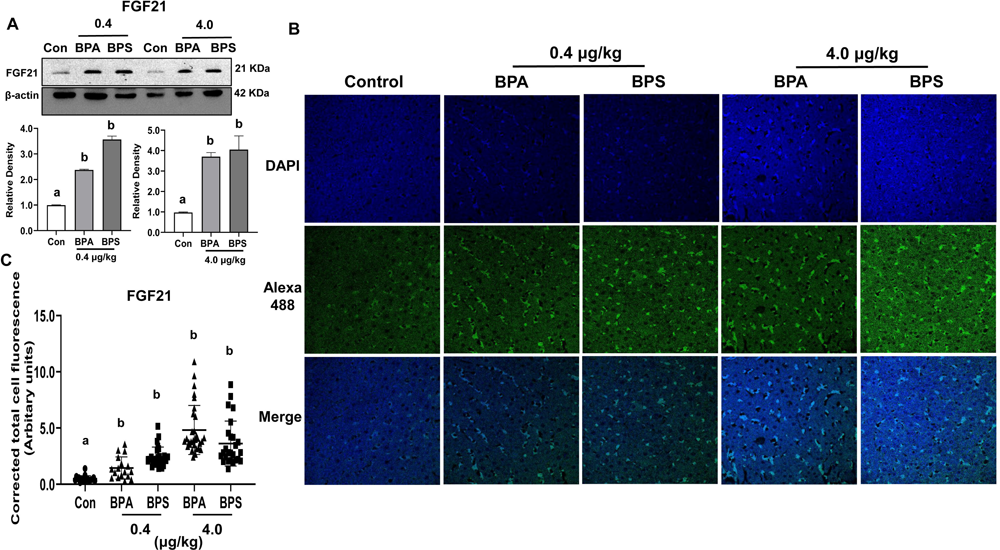
The expression and localization of a liver-secreted peptide hormone, FGF21, in the 90d offspring liver tissues following bisphenols exposure prenatally. (A) Immunoblots of FGF21 and its relative expression (n=3 rats/group). (B) Localized expression of FGF21 protein by immunofluorescence as described in Fig.6, where Alexa Fluor 488 tagged FGF21 antibody (green) and nuclei with DAPI (blue). (C) The total fluorescence of FGF21 was quantified as mentioned (n=3). Data were analysed with one-way ANOVA with Tukey’s multiple comparison tests and represented as means ± SEM. Values with unlike letters were significantly different vs control at p<0.05.

### 3.8 Prenatal bisphenol exposure induced liver inflammation by increased pro-inflammatory cytokines expression and oxidative stress in the offspring

BPS-exposed offspring liver increased the expression of IL6, CRP, IL1β, TNFα, and COX2, indicating chronic inflammasome activation (**Fig.7 A-E**). The MDA levels were consistently increased in the BPS-exposed liver across the concentration [control vs. BPS (0.4 μg/kg) and BPS (4.0 μg/kg): 1.58±0.19 vs. 4.12±0.99 and 4.77±0.86 (nmol/mg), p < 0.05 **(Fig.7F)**] indicating deterioration of the lipids membrane of the liver leading to disintegration and dysfunctions at the cellular and sub-cellular levels.

**Fig.7.**
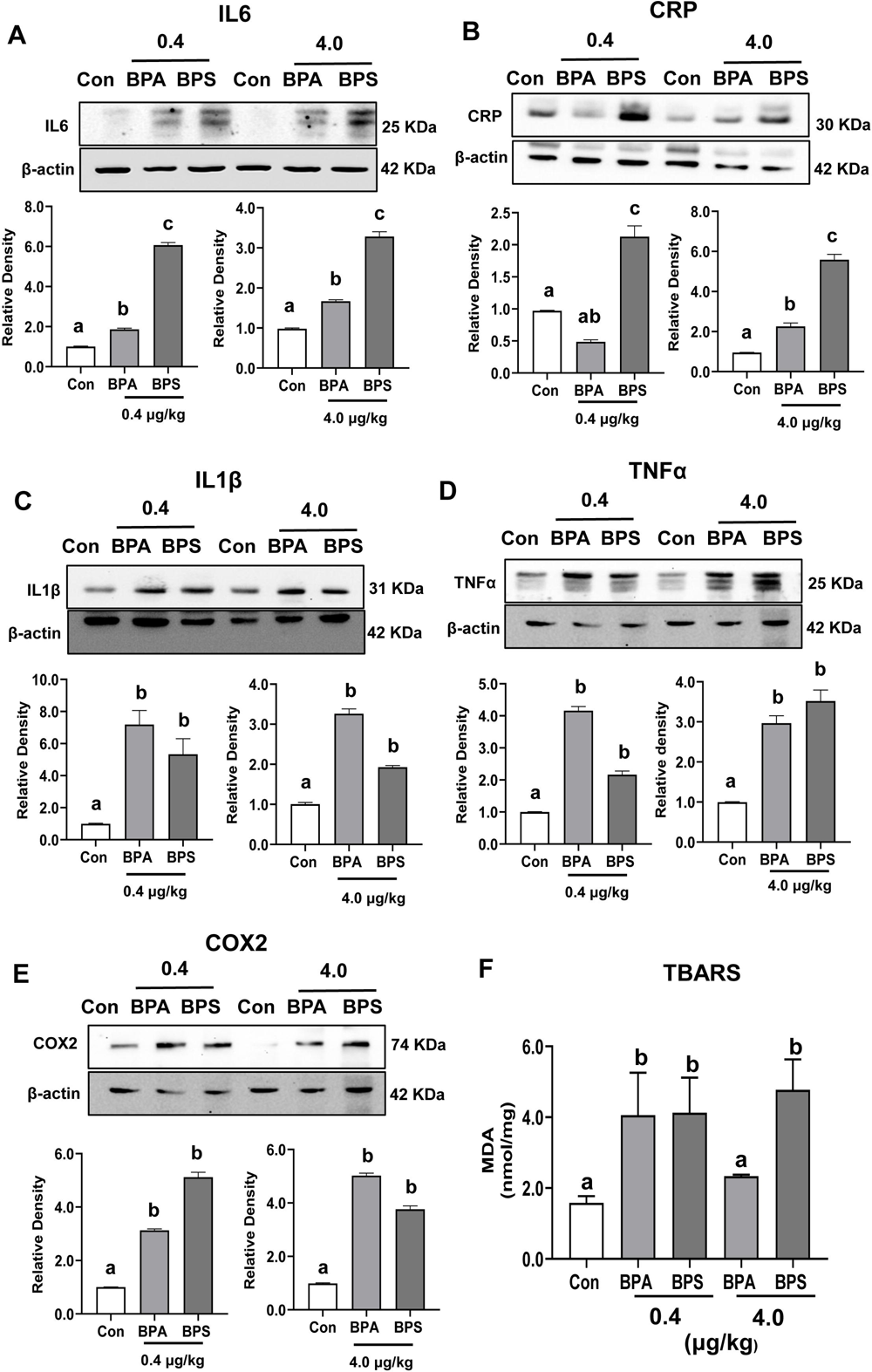
Expression of pro-inflammatory cytokines and lipid peroxidation in the offspring’s liver tissue following prenatal bisphenol exposure. (A-E) Immunoblots and their respective relative band intensities of (A) IL6, (B) IL1β, (C) CRP, (D) TNFα, and (E) COX2 are shown. Respective bars indicate the relative band intensities of normalized expressed proteins (n=3 rats/group). (F) Levels of malondialdehyde (MDA) produced as an end product of thiobarbituric acid reactive substances (TBARS) were measured in liver lysates (n=6-8/group). Data represent mean ± SEM using one-way ANOVA with Tukey’s multiple comparison test. Values with unlike letters indicate significantly different at p<0.05.

## 4. Discussion

This study, for the first time, demonstrated that fetal exposures to BPS exhibited a more significant effect than BPA in promoting adipocyte hypertrophy, dyslipidemia, oxidative stress, and microvesicular steatosis by activating the glucocorticoid system in the offspring fed with a standard chow diet. Liver histology indicated excess lipid droplets, hepatocyte ballooning, a higher ADRP expression in the BPS-exposed offspring, and developed obese phenotype like BPA, followed by increased expression of pro-inflammatory cytokines in adipose and liver. Overweight and pale livers were elevated with lipid load (triacylglycerol and cholesterol), inflated hepatocytes knitted with numerous lipid droplets, upregulated expression *de novo* lipogenesis, and glucocorticoid mediators. The bisphenol-exposed fatty liver in the offspring could arise from excess adipose lipid uptake and inflammation. Moreover, prenatal BPS exposure predisposed to steatosis in adult life to a greater extent than those exposed to BPA.

The LEPR mRNA overexpression promotes diet-induced obesity in rodents (Gamber et al., 2012), indicating that leptin resistance might have resulted in higher feed efficiency and obesity in BPS-exposed offspring. Unlike gestational BPS-exposed male mice in HFD (Ivry Del Moral et al., 2016), male rat offspring fed a regular chow diet developed obesity in our study, indicating the endocrine-dependent effects predominated over calorie intake. The BPS-exposed offspring showed an association of giant white adipose tissue fat cells with higher body fat, considered the centre of metabolic dysfunction.

It is suggested that adipocyte hypertrophy contributes to liver fat accumulation independent of other causes (Petäjä et al., 2013). NAFLD patients had larger adipocytes than healthy, which is involved in the pathophysiology of hepatic steatosis (Holmer et al., 2022), indicating a dysregulation of adipose tissue metabolism due to altered adipocyte size and function. Prenatal BPS exposure induced expansion of adipocyte size (60-90 µm) with larger fat cells than control adipocytes (40-50 µm). In the non-obese state, adipocyte size distribution is highly homogeneous, and the diameter of the large adipose cells was associated with a higher body fat percentage (McLaughlin et al., 2014). The BPS exposure dysregulated both of this homeostasis in adipose tissue. The increased adipocyte size reflects abnormal FFA uptake, lipid droplet triacylglycerol storage, and also FFA release by lipolysis.

Moreover, several mediators that regulate lipid homeostasis might control adipocyte size. The reduced PPARγ and increased FABP4 expression in BPS-exposed adipose tissue can regulate the adipocyte size independent of insulin resistance. BPS binds PPARγ with similar efficacy as its natural ligand (GW9662) to modulate pre-adipocyte differentiation and metabolic signalling (Schaffert et al., 2021). This effect, mediated through inhibition of PPARγ, could promote BPS-induced adipocyte hypertrophy. Increased adipocyte size in PPARγ knock-out mice (Gumbilai et al., 2016) and its low expression in mature adipocytes (Sauma et al., 2007) indicate that PPARγ regulates their size. Thus, decreased PPARγ expression in BPS-exposed adipose might reduce PPARγ activity to facilitate lipid uptake by increasing adipocyte size. Again, increased FABP4 negatively correlates with PPARγ in adipose tissue as FABP4 induces proteasomal degradation of PPARγ (Garin-Shkolnik et al., 2014). The enlarged adipocyte might be favoured by FABP4 accumulation due to its increased expression in adipose, reducing PPARγ, and AMPK activity, leading to increased basal lipolysis (Berger and Géloën, 2023). Since large adipocytes have a two-fold higher basal lipolysis rate than the small ones (Wueest et al., 2009), BPS exposure might have induced a higher basal lipolysis rate due excess presence of large adipocytes. The obesity-induced adipocyte expansion might involve adipose tissue remodelling by triggering apoptosis and ER stress (Kawasaki et al., 2012). The unfolded proteins accumulate in the ER lumen, leading to programmed cell death. Upregulated CHOP and effector caspase 3 expression indicate many apoptotic adipocytes in bisphenol-exposed adipose tissue.

Adipocyte hypertrophy relates to several factors, including impaired adipose tissue differentiation, rate of adipogenesis, adipocyte senescence and necrosis, and adipose tissue inflammation. Adipocyte hypertrophy results in inflammation in adipose tissue and secretion of chemokines that might have attracted neutrophil-infiltrated immune cells in the development of obesity (Mohammed et al., 2022). As evidenced in BPS-exposed adipocytes, neutrophil infiltration could enhance adipose tissue inflammation. In an obese state, adipose tissue macrophages secrete many pro-inflammatory cytokines that can induce inflammation. The metaflammation (low-grade chronic adipose tissue inflammation) is consistently associated with excess body fats, adiposity, hyper corticosterone, and expression of pro-inflammatory cytokines observed in BPS-exposed offspring, indicating severely inflamed adipose tissue. The macrophage numbers might have elevated in bisphenol-exposed adipose tissues as their increased accumulation activated the pro-inflammatory cascade (Ampem et al., 2019). BPS-exposed rats become obese due to severe metabolic and systemic inflammation impairment propelled by macrophage accumulation (Weisberg et al., 2003), as evidenced by upregulated IL1β, CRP, and COX2 expression in our study. Adipose tissue macrophages are rich in cyclooxygenase (COX), a rate-limiting enzyme for producing prostaglandin E2 which increases adipose tissue inflammation. The rapid surge in COX2 expression over 20 to 30-fold in BPS-exposed VAT might play a compensatory protective role in lowering adipose tissue dysfunction (Pan et al., 2022) and delaying NAFLD progression (Motiño et al., 2016). BPS-induced adipocyte hypertrophy could be due to ectopic fat deposition promoting increased IL1β expression since its upregulated expression releases the fatty acids into the circulation during obesity (Bing, 2015).

The offspring exposed to BPS showed an elevated corticosterone in plasma and increased expression of HSD11β1 in adipose tissue than control indicating impaired glucocorticoid metabolism. The increased metabolic clearance of corticosterone occurs due to increased physiological distribution of corticosterone and/or altered HSD11β1 activities responsible for corticosterone metabolism in obese rats (White et al., 1989). Unlike obese rats, the metabolic clearance rate of BPS-exposed rats did not correlate with body weight. However, overexpression of HSD11β1 might be responsible for increased corticosterone concentrations. In addition, elevated serum cortisol was correlated with the histopathological severity of the NAFLD (Targher et al., 2006), indicating a risk to the progression of NAFLD of the BPS-exposed offspring due to their closer histopathology and elevated corticosterone.

Prenatal BPS exposure might have reprogrammed the normal glucocorticoid metabolism in the offspring. The overexpressed HSD11β1 in the VAT and elevated corticosterone indicated their roles in the development of steatosis. In humans, increased HSD11β1 expression in VAT was associated with excess adipogenesis and NAFLD (Candia et al., 2012). Adipogenesis is governed by the action HSD11β1, which converts inactive cortisone into active corticosterone and activates the glucocorticoid receptors (GC). The deleterious metabolic effects of visceral obesity are associated with the activation of GC by HSD11β1 (Pereira et al., 2012). Although BPS-exposed rats did not affect glucose levels, they developed obesity with excess visceral adiposity at their adult (90d) age. Elevated expression of HSD11β1 causes an imbalance of cortisol and glucocorticoid levels that might result in metabolic imbalance, impaired clearance of hepatic lipids, and develop dyslipidemia, leading to hepatic steatosis.

The elevated cholesterol and triacylglycerol, oxidative stress, and upregulation of PPARα of the BPS-exposed offspring liver suggest that increased accumulation of FFAs contributed to liver weight and discoloration. Lipid deposition, an excess gross size, and the pale yellowish appearance of the liver surface are signs of steatosis. The excessive triacylglycerol accumulation characterizes hepatic steatosis originating from the adipose tissue and endocytosis remnants of triacylglycerol-rich lipoproteins. In addition*, de novo* lipogenesis triggers VLDL secretion, and the redundant release of FFAs from AT triggers hepatic steatosis (Jiang et al., 2016). Steatosis involves excess lipid acquisition over its disposal due to higher hepatic lipid uptake, *de novo* lipogenesis, oxidative stress, and mitochondrial dysfunctions (Ipsen et al., 2018). Several of these features were aligned with BPS-exposed offspring.

Due to its limited storage capacity, the efflux of excess FFA from adipose tissue results in ectopic fat accumulation in the liver by enhancing *de novo* lipogenesis; the latter is increased in the NAFLD (Lambert et al., 2014). Since total triacylglycerol and cholesterol are key parameters to confirm the onset of the NAFLD risk (Fukuda et al., 2016), their elevation in BPS-exposed offspring predisposes NAFLD risk. In our study, dyslipidemia due to BPS exposure was similar to HFD-induced obesity (Labaronne et al., 2017). Inflammation is often associated with elevated triacylglycerol in obesity, where hypertrophied adipocytes secrete chemokines, promoting macrophage infiltration in adipose tissue, thereby contributing to a pro-inflammatory response (Zhang et al., 2018). Thus, BPS exposure mimics the conditions that a HFD has established.

The mechanism of BPS-induced hepatic lipid deposits probably results from the increased *de novo* lipogenesis promoted by upregulated expression of sterol and lipid biogenesis (SREBF1, FASN, ACACA, NAMPT) and glucocorticoid genes. Fetal BPS exposure might destabilize the homeostasis of liver metabolic genes expression by epigenetic programming since the obesogenic effect of BPS was mediated by hypomethylation of a specific set of gene promoters involved in lipid metabolism and fibrosis in the liver (Brulport et al., 2020).

The BPS-exposed liver showed larger lipid droplets, hepatocyte ballooning vis-à-vis much higher expression of the ADRP, and its localization in the hepatocyte than BPA. Steatosis, a part of NAFLD, is observed as a large number of lipid droplets in which the hepatocyte is filled with fat, displacing the nucleus to the periphery. Microvesicular steatosis is characterized by hepatocyte ballooning and possibly impaired mitochondrial β-oxidation, making it vulnerable to the advanced state of NAFLD (Tandra et al., 2011). Although we did not measure fatty acid oxidation, the dysregulated expression of ADRP in the liver and their increased abundance in hepatocytes might involve BPS-induced microvesicular steatosis. ADRP is instinctively associated with lipid droplets on their surface, its abundance is proportionate to the amount of intracellular lipid content, and its involvement is indispensable in the development of the fatty liver. Upregulated ADRP expression in mice models of steatosis fed with a HFD (Motomura et al., 2006), suggest a comparable effect in BPS-exposed offspring could lead to the progression of NAFLD by ADRP overexpression. ADRP expression is transcriptionally regulated by PPAR-responsive elements that control the transcription of the ADRP gene in hepatocytes (Targett-Adams et al., 2005). PPARα upregulation in BPS-exposed liver might involve a compensatory protective enhancement of fatty acid oxidation in normalizing the elevated lipid levels.

An increased FGF21 expression in the liver and their extensive localization in hepatocytes might play a role in the BPS-induced microvesicular steatosis. FGF21 is a hepatokine predominantly produced by the liver and released into the circulations, where its level is increased in obesity, hypertriglyceridemia, and impaired glucose tolerance. The FGF21 level is increased at liver injury sites in NAFLD (Dushay et al., 2010), as a compensatory mechanism to protect the liver from ER stress, lipotoxicity, and dysregulated metabolism (Rusli et al., 2016). The circulatory FGF21 and its expression in the liver are elevated in hepatitis subjects, a biomarker to detect the early stages of NAFLD (Yang et al., 2013). The upregulated FGF21 expression and its localization in BPS-exposed liver could be proportional to the progression of steatosis, providing an extenuatory effect to maintain lipid homeostasis.

Fetal exposure to BPS activated inflammatory cascades at the systemic and tissue level in the offspring by increased expression of several pro-inflammatory mediators such as C-reactive protein (CRP), IL1β, IL6, COX2, and TNFα in offspring adipose and liver tissues. IL1β is activated by damage-associated molecular pattern together with TLR ligands, further activates NLRP3 and NLRP6 inflammasomes, thereby inducing caspase-1 activation and releasing IL1β in hepatocytes, promoting low-grade inflammation in the liver (Barbier et al., 2019). Hepatic macrophage dysfunction leads to the development of steatohepatitis due to overexpressing TNFα and IL6 in obesity-linked NAFLD (Kanda et al., 2020). On BPS exposure, the offspring’s liver was prone to oxidative lipid peroxidation, cell death, and endoplasmic reticulum stress, collectively leading to hepatic steatosis.

The study has several limitations, including that it lacks analysis of fatty liver by oil-o-red staining. Moreover, we could not measure mitochondrial oxidation to measure lipid metabolism in the offspring.

In several ways, BPS mediated its metabolic effects differently than BPA. BPS induced adipocyte hypertrophy more than BPA due to differential preadipocyte differentiation. Unlike BPA, which induces adipogenic differentiation potential gene expression, BPS affected adipogenic terminal differentiation expression in male preadipocytes (Pu et al., 2017). During differentiation, committed preadipocytes undergo growth arrest and subsequent terminal differentiation into mature adipocytes, which is accompanied by a dramatic surge in the expression of adipocyte fatty acid binding protein and lipid-metabolizing enzymes as evidenced in BPS-exposed offspring adipose tissue in our study. The BPS-exposed liver gained a higher weight and more discoloration, elevated total cholesterol and triacylglycerol, and oxidative stress than BPA at the respective concentration. Like BPA, BPS also interacts with PPARγ, a pivotal regulator of adipogenesis with similar efficiency (Schaffert et al., 2021). BPS systemic clearance is two-fold slower than BPA, with an average elimination half-life of 7.9-9.3 h of the labelled (deuterated) BPS and BPS glucuronide in humans (Khmiri et al., 2020). The differences in the kinetics indicate much higher systemic levels of active BPS than BPA and predict a higher oral bioavailability of BPS than BPA via all possible exposure routes.

## 5. Conclusions

Although BPA-containing products are declining from consumer merchandise, the supply of BPA-free products has gained huge consideration globally. Accumulating evidence substantiates subtle exposure risks for developing metabolic diseases and questions the safety of life-course exposure to BPS for health and disease risks. BPS exposure is causally implicated with the association of metabolic disease risk factors. Considering these, European Food Safety Authority (EFSA) made regulations on tolerable daily intake (TDI) for BPS 0.05 mg/kg bw/day, a value agreed upon by other regulatory bodies, including the U.S. Environmental Protection Agency (1993). Our data suggest that less than 100-fold of TDI level exposure of BPS (0.4 µg/kg bw/day) to pregnant dams severely affected liver and adipose metabolic regulation in the 90d male offspring fed a standard chow diet. However, similar exposure of this BPA-alternative chemical on human metabolic functions is required to confirm these risks.

## Authors’ contributions

AM conducted the animal trial, sample and data collection, performed major laboratory experiments, data analysis and wrote the draft; SV conducted the animal trial, sample, and data collection and performed key laboratory experiments, analysis, and data presentation. KSN, MS, and SRK performed the experiments. AI commented on the draft. AKD provided critical comments and reviewed the manuscript. SB fund acquisition & conceptualized, supervised, administered, drafted, interpreted, and finalized the manuscript.

