## Supplementary material for "Gestational exposure to bisphenol S induces microvesicular steatosis by promoting lipogenesis and inflammation in male rat offspring": Sup table 1

| **Target**  **protein** | **Protein**  **name** | **Primary antibody** | **Clonality** | **Host** | **Catalog & supplier** | **Dilution** |
| --- | --- | --- | --- | --- | --- | --- |
| TNFα | Tumor necrosis factor | Anti-TNFα | Polyclonal | Rabbit | #3707s CST | 1:1000 |
| IL6 | Interleukin 6 | Anti-IL6 | Polyclonal | Rabbit | #P160 Thermo | 1:1000 |
| IL1β | Interleukin 1 beta | Anti- IL1β | Monoclonal | Mouse | #sc-52012 Santa Cruz | 1:1000 |
| NFκB | Nuclear factor kappa B subunit 1 | Anti- NFκB | Monoclonal | Mouse | #sc-8008 Santa Cruz | 1:1000 |
| CRP | C-reactive protein | Anti-CRP | Monoclonal | Mouse | #sc-69770 Santa Cruz | 1:1000 |
| COX2 | Prostaglandin-endoperoxide synthase 2 | Anti-COX2 | Polyclonal | Rabbit | #PA5-32366 Thermo | 1:3000 |
| 11β-HSD1 | Hydroxysteroid 11-beta dehydrogenase 1 | Anti-11β-HSD1 | Polyclonal | Rabbit | #sc-20175 Santa Cruz | 1:1000 |
| CHOP | DNA damage inducible transcript 3 | Anti-CHOP | Monoclonal | Mouse | #2895 CST | 1:1000 |
| Caspase 3 | Cystine-aspartic acid protease | Anti-Caspase 3 | Polyclonal | Rabbit | #PA5-77887 - Thermo | 1:2000 |
| FGF21 | Fibroblast growth factor 21 | Anti-FGF-21 | Monoclonal | Rabbit | #a3908-ABclonal | 1:1000 |
| FABP4 | Fatty acid binding protein 4 | Anti-FABP4 | Polyclonal | Rabbit | #PA5-30591 Thermo | 1:2500 |
| ADRP | Perilipin 2 | Anti-ADRP | Polyclonal | Rabbit | #PA5-29099 Thermo | 1: 1000 |
| PPARα | Peroxisome proliferator activated receptor alpha | Anti-PPAR α | Monoclonal | Mouse | #MA1-822 Thermo | 1:1000 |
| PPARγ | Peroxisome proliferator activated receptor gamma | Anti-PPAR γ | Monoclonal | Mouse | #sc-7273 Santa Cruz | 1:500 |
| Actin | Beta actin | Anti-β Actin | Monoclonal | Mouse | #A5316 Sigma | 1:10000 |

**Table S1: The primary antibodies and their dilution used in this study**
