## Supplementary material for "Gestational exposure to bisphenol S induces microvesicular steatosis by promoting lipogenesis and inflammation in male rat offspring": Sup table 2

**Table S2**: Predesigned SYBR green I rat primers and corresponding genes used for the mRNA expression analyses

| **Sl.**  **no.** | **Primer**  **ID** | **Gene symbol** | **Gene ID** | **Gene name** | **Nucleotide sequences (5’-3’)** | **Ref_seqID** |
| --- | --- | --- | --- | --- | --- | --- |
| 1 | R1_Esr1 | ESR1 | [24890](https://www.ncbi.nlm.nih.gov/gene/24890) | Estrogen receptor 1 | F 5'- ATATGATCAACTGGGCAAAG-3'  R 5'- CATTTACCTTGATTCCTGTCC-3' | [NM_012689](http://www.ncbi.nlm.nih.gov/entrez/query.fcgi?db=Nucleotide&cmd=Search&term=NM_012689&doptcmdl=GenBank) |
| 2 | R1_Ar | AR | [24208](https://www.ncbi.nlm.nih.gov/gene/24208) | Androgen Receptor | F 5'- CCTTGTTCCCTTTTCAGATG-3'  R 5'- GTAAAAGAGGCAGAGAAGAAG-3' | [NM_012502](http://www.ncbi.nlm.nih.gov/entrez/query.fcgi?db=Nucleotide&cmd=Search&term=NM_012502&doptcmdl=GenBank) |
| 3 | R1_Hsd11β1 | HSD11β1 | [25116](https://www.ncbi.nlm.nih.gov/gene/25116) | 11beta-hydroxysteroid dehydrogenase type 1 | F 5'- TAGACACAGAAACAGCTTTG-3'  R 5'- AATTCCATGATCCTCCTTCC-3' | [NM_017080](http://www.ncbi.nlm.nih.gov/entrez/query.fcgi?db=Nucleotide&cmd=Search&term=NM_017080&doptcmdl=GenBank) |
| 4 | R1_Hsd11β2 | HSD11β2 | [25117](https://www.ncbi.nlm.nih.gov/gene/25117) | 11beta-hydroxysteroid dehydrogenase type 2 | F 5'- CAGGAGACATGCCATACC-3'  R 5'- GATGATGCTGACCTTGATAC-3' | [NM_017081](http://www.ncbi.nlm.nih.gov/entrez/query.fcgi?db=Nucleotide&cmd=Search&term=NM_017081&doptcmdl=GenBank) |
| 5 | R1_Hsd3β1 | HSD3β1 | [360348](https://www.ncbi.nlm.nih.gov/gene/360348) | 3β-hydroxysteroid dehydrogenases | F 5'- CCAAGGTGACAATGTTAGAAG-3'  R 5'- TATGTTCTGGGTACCTTTCAG-3' | [NM_001007719](http://www.ncbi.nlm.nih.gov/entrez/query.fcgi?db=Nucleotide&cmd=Search&term=NM_001007719&doptcmdl=GenBank) |
| 6 | R1_Igf1 | IGF1 | 24482 | Insulin like growth factor 1 | F 5'- GCACCTCCAATAAAGATACAC-3'  R 5'- TGGGCTTGTTGAAGTAAAAG-3' | NM_001082479 |
| 7 | R1_Fabp4 | FABP4 | 79451 | Fatty acid-binding protein 4 | F 5'- AAGTGAAGAGCATCATAACC-3'  R 5'- TGATGCAAATTTCAGTCCAG-3' | [NM_053365](http://www.ncbi.nlm.nih.gov/entrez/query.fcgi?db=Nucleotide&cmd=Search&term=NM_053365&doptcmdl=GenBank) |
| 8 | R1_Slc2a4 | GLUT4 | [25139](https://www.ncbi.nlm.nih.gov/gene/25139) | Glucose transporter member 4 | F 5'- AAGTGATTGAACAGAGCTAC-3'  R 5'- CTTTTCCTTCCCAACCATTG-3' | [NM_012751](http://www.ncbi.nlm.nih.gov/entrez/query.fcgi?db=Nucleotide&cmd=Search&term=NM_012751&doptcmdl=GenBank) |
| 9 | R1_Scd-1 | SCD 1 | 246074 | Stearoyl-Coenzyme A desturase 1 | F 5'- ATGAGAGAAGATATCCACGAC-3'  R 5'- AGTAAAATATCCCCCAGAGC-3' | NM_139192.2 |
| 10 | R1_Fasn | FASN | 50671 | Fatty acid synthase | F 5'- AAAAGGAAAGTAGAGTGTGC-3'  R 5'- GACACATTCTGTTCACTACAG-3' | NM_017332.2 |
| 11 | R1_Srebp1 | SREBP1 | 78968 | Sterol regulatory element binding transcription factor 1 | F 5'- AAACCTGAAGTGGTAGAAAC-3'  R 5'- TTATCCTCAAAGGCTGGG-3' | NM_001276707 |
| 12 | R1_Cebpα | CEBPα | 24252 | CCAAT/enhancer binding protein alpha | F 5'- AAGAGCCGAGATAAAGCC-3'  R 5'- GTCATTGTCACTGGTCAAC-3' | NM_001287579 |
| 13 | R1_Nampt | NAMPT | 297508 | Nicotinamide phosphoribosyltransferase | F 5'- GAATATGGCCATGATCTTCTC-3'  R 5'- TTTTTCTGACTTCGTCAAACG-3' | NM_177928 |
| 14 | R1_Prkaa2 | PRKAA2 | 78975 | Protein kinase AMP-activated catalytic subunit alpha 2 | F 5'- GAATGGAAGGTAGTGAATGC-3'  R 5'- TAAAGTCTAGAAGATAGCTCCG-3' | NM_023991 |
| 15 | R1_Acaca | ACACA | 60581 | Acetyl-CoA carboxylase alpha | F 5'- AGCAGTATTTGAACACATGG-3'  R 5'- CAGTTCCAAGAAGTAGAAGC-3' | NM_022193 |
| 16 | R1_Ppar-α | PPAR-α | 25747 | Peroxisome proliferator activated receptor alpha | F 5'- CTGCTATAATTTCGTGTGGAG-3'  R 5'- GAGTTTTGGGAAGAGAAAGG-3' | NM_013196.2 |
| 17 | R1_Ppar-γ | PPAR-γ | 25664 | Peroxisome proliferator activated receptor gamma | F 5'- AAGACAACAGACAAATCACC-3'  R 5'- CAGGGATATTTTTGGCATACTC-3' | NM_013124.3 |
| 18 | R1_Lep | LEP | 25608 | Leptin | F 5'- CTCATCAAGACCATTGTCAC-3'  R 5'- TGAGGATCTGTTGATAGACTG-3' | NM_013076.3 |
| 19 | R1_LepR | LEPR | 24536 | Leptin receptor | F 5'- TGTAAAAGTTCCTATGAGAGGG-3'  R 5'- CATACCTCCTCACACTACAC-3' | NM_012596 |
| 20 | R1_Adipoq | ADIPOQ | 246253 | Adiponectin | F 5'- TGGCGATTTTCTCTTCATTC-3'  R 5'- AGGATTAAGAGGAACAGGAG-3' | NM_144744.3 |
| 21 | R1_Pomc | POMC | 24664 | Proopiomelanocortin | F 5'- AACGCCATCATCAAGAAC-3'  R 5'- AAGGTTTTATTTCCTAACTACAG-3' | NM_139326 |
| 22 | R1_Npy | NPY | 24604 | Neuropeptide Y | F 5'- AGACAGAGATATGGCAAGAG-3'  R 5'- TTCACAGGATGAGATGAGATG-3' | NM_012614 |
| 23 | R1_Fads1 | FADS1 | 84575 | Fatty acid destaurase 1 | F 5'- GTACTTCTTCTTGATTGGAC-3'  R 5'- GTAAGTGAAGAAGACACGAAC-3' | NM_053445.2 |
| 24 | R1_Fads2 | FADS2 | 83512 | Fatty acid desaturase 2 | F 5'- CTTCTTCAATGACTGGTTCAG-3'  R 5'- CTTCAGTGAACTCACAATGTC-3' | NM_031344.2 |
| 25 | R1_Elovl2 | ELOVL2 | 498728 | Fatty acid elongase 2 | F 5'- CTTGTGGTCAAAGCTTCTTC-3'  R 5'- GAGGTATTTCTTCCACCAAAG-3' | NM_001109118.1 |
| 26 | R1_Elovl5 | ELOVL5 | 171400 | Fatty acid elongase 5 | F 5'- TTCTTCGTAAGAACAACCAC-3'  R 5'- ATAGTACGAGTACATGAGGAC-3' | NM_134382.2 |
| 27 | R1_Act-B | ACT β | 81822 | Actin-beta | F 5'- AAGACCTCTATGCCAACAC-3'  R 5'- TGATCTTCATGGTGCTAGG-3' | NM_031144.3 |
