## Supplementary material for "Gestational exposure to bisphenol S induces microvesicular steatosis by promoting lipogenesis and inflammation in male rat offspring": sup table 3

**Table 1.** Effects of gestational bisphenols exposure on biochemical indices in the plasma parameter of the 90 days male offspring rats **^#^**

|  | **Experimental groups (µg/kg bw)** | | | | |
| --- | --- | --- | --- | --- | --- |
| **Parameters** | **Con (0.0)** | **BPA (0.4)** | **BPS (0.4)** | **BPA (4.0)** | **BPS (4.0)** |
| Glu (mg/dl) | 118.3±2.58 | 116.3±3.21 | 119.1±5.53 | 112.8±2.08 | 115.6±4.01 |
| Insulin |  |  |  |  |  |
| TG (mg/dl) | 140.8±3.0 | 149.5±6.42 | 153.0±7.33 | 135.7±3.16 | 140.1±3.27 |
| TC (mg/dl) | 112.3±1.15 | 109.5±3.60 | 98.79±6.83 | 105.0±3.21 | 107.4±7.25 |
| Cort (ng/ml) | 12.19±0.88 | 30.20±3.28* | 37.08±6.46* | 15.71±2.57 | 25.92±4.18 |

^#^ Data were analysed with one-way ANOVA with post-hoc Sidak’s multiple comparison test. Values are represented as means ± SEM (n=6-8/group). * p<0.05 was considered as statistically significant vs. control. Glu: Fasting glucose; TG: Total triglycerides; TC: Total cholesterol; Cort : Corticosterone.
